# Haploinsufficiency of KPNA7 causes otosclerosis, likely due to the release of import inhibition of PTHrP and the reactivation of chondrogenesis in the globuli interossei

**DOI:** 10.1101/2025.08.22.671726

**Authors:** Tammy Benteau, Nelly Abdelfatah, Anne Griffin, Cindy Penney, Pingzhao Hu, Susan G. Stanton, Guangju Zhai, Maxime Maheu, Terry-Lynn Young

## Abstract

Otosclerosis is a genetic bone disorder restricted to the otic capsule and a common cause of conductive hearing loss with both familial and sporadic cases. To date, 14 genomic loci (*OTSC*) and four underlying *OTSC* genes (*MEPE*, *SERPINF1, FOXL1, SMARCA4)* have been identified in autosomal dominant families. A combined genetic/genomics approach on five affected siblings of Northern European ancestry from the island of Newfoundland, Canada identified a premature stop mutation in Karyopherin subunit α7 (*KPNA7*, c.49C>T, p.R17X). *KPNA7* maps to *OTSC2* (7q22.1) and encodes the newest of the seven-member importin-α family of nuclear transporters and plays a critical role in early embryonic cleavage events and zygotic genome activation. Previous studies reveal that recessive *KPNA7* variants cause skeletal abnormalities, including scoliosis and ocular hypertelorism in two sisters with Partial Corpus Callosum Agenesis-Cerebellar Vermis Hypoplasia With Posterior Fosa Cysts Syndrome and more recently, have been implicated in preimplantation embryo arrest (PREMBA) (OMIM 614107). Interestingly, *KPNA7* is also a maternal factor with an exclusively embryonic role and likely inhibits non-classical NLS transport of PTHrP, a known activator of chondrogenesis. We propose that *KPNA7* haploinsufficiency causes a failure in nuclear transport inhibition of PTHrP in the quiescent embryonic cells of the globuli interossei in the otic capsule and re-activates chondrogenesis. The KPNA7 discovery provides new insights into the pathogenesis of otosclerosis and potential for targeted therapies.

**Author Summary:** Otosclerosis is a distinctly human genetic bone disorder of the otic capsule and a major cause of progressive hearing loss in young adults, particularly in females. Even though otosclerosis has been recognized as a distinct entity for a long time, both its pathogenesis and restriction to the otic capsule remains a mystery. Here, we use a combined genetic/genomics approach to identify a premature stop mutation in five affected siblings of Northern European ancestry from the island of Newfoundland, Canada. KPNA7 encodes the newest of the seven-member importin-α family of nuclear transporters and plays a critical role in early embryonic cleavage events and zygotic genome activation. Based on the unique features of the otic capsule, we hypothesize that the premature stop mutation in *KPNA7* leads to haploinsufficiency causing a failure in nuclear transport inhibition of PTHrP and reactivates chondrogenesis in the otherwise quiescent embryonic cells within the otic capsule. The KPNA7 discovery provides new insights into the pathogenesis of otosclerosis and potential for targeted therapies.

## Introduction

Otosclerosis is a uniquely human skeletal disorder restricted to the otic capsule of the temporal bone and a common cause of progressive conductive hearing loss (HL) in young adults. An autosomal dominant (AD) disease with environmental triggers, otosclerosis is clinically characterized by abnormal bone deposition in the middle ear, distorting the fine structures of the ossicular chain and limiting the movement of the stapes bone against the oval window. The immobilization of the stapes bone results in conductive HL as well as sensorineural HL in some patients due to abnormal bone growth extending into the fluid-filled inner ear (1). The location of the inner ear within the osseous labyrinth greatly impacts perilymph sampling for diagnostic purposes and local drug delivery. Visualization during surgery to replace or repair stapes fixation due to otosclerosis validates the diagnosis, and in most cases, restores the conductive component of hearing to those able to access stapes surgical prosthesis replacement. Otosclerosis risk factors include positive family history, sex (female), measles and pregnancy (1, 2). Recognised as a medical entity some 125 years ago (3), this restricted bony disorder cannot be predicted, stopped or medically treated, and its pathogenesis remains a mystery.

Despite several decades of research efforts, the otosclerosis (*OTSC*) genes have been recalcitrant to discovery because of the genetically heterogeneous nature of otosclerosis and the rarity of AD families under study (2). So far, 14 distinct (*OTSC1-14)* loci have been mapped in AD families and four causative *OTSC* genes (*MEPE*, *SERPINF1, FOXL1, SMARCA4)* identified (Hereditary Hearing Loss Homepage). A recent search for susceptibility factors involving 3504 otosclerosis cases from three biobank studies revealed 23 novel loci linked to genes whose dysregulation in bone remodeling and mineralization causes rare monogenic skeletal disorders (4). These near protein associations provide insight into the nature of the highly penetrant *OTSC* genes but not their identity, as GWAS studies exclude rare variants by design. Given the high genetic heterogeneity underlying skeletal dysplasias, with 461 genes known to cause monogenic forms, more *OTSC* genes are anticipated (5).

The otic capsule has a highly complex anatomy, and its embryonic development is one of the most complicated examples of cellular morphogenesis in any biologic system (6). In temporal bone, the inner ear tissues and spaces are enclosed within the bony otic capsule, the hardest bone in the body, a critical feature essential to maintaining hearing integrity (7). The otic capsule forms through endochondral ossification, one of two essential pathways of bone formation that uses cartilage as a bone template during fetal development. Mesenchymal stem cells differentiate into chondrocytes (cartilage cells) which proliferate rapidly, hypertrophy and secrete the extracellular matrix that undergoes mineralization. Eventually the hypertrophic chondrocytes die through apoptosis and are replaced by osteocytes that become trapped in bony matrix. Although the otic capsule is fully formed by the fifth fetal month (8), islands of embryonic tissue containing quiescent chondrocytes and osteocytes, known as the globuli interossei, are uniquely retained by the otic capsule and persist throughout life. These embryonic remnants are increasingly implicated as the site of otosclerosis in the temporal bone (9–11).

The human skeleton continues to grow and repair (remodel) postnatally via the well-studied RANK/RANKL/OPG pathway. In contrast, the otic capsule and the ossicular chain are fully formed *in utero* and remodeling is virtually absent due to the overproduction of OPG (osteoprotegerin) (12). In the bony lacunae of the otic capsule, osteocytes communicate via an intercellular canalicular network that provides nutrient transport and bathes the perilacunar matrix with OPG. As a normal part of aging, osteocytes die but are not replaced as OPG prevents osteoclasts from maturing and beginning the remodeling process (13). How bone remodeling in the otic capsule occurs in the presence of overproduction of OPG close to the inner ear is a mystery, and suggests other parallel protective mechanisms must play a role (13, 14).

Herein we identify a stop mutation in *KPNA7* (Karyopherin subunit α7), the newest member of the Importin-α (Imp-α) transport factors involved in nucleocytoplasmic trafficking, in a white family of Northern European descent. Nucleocytoplasmic trafficking is a highly efficient and regulated system comprising of 60 proteins of the nuclear transport system where dysregulation is linked to major diseases such as cancer, viral infections, inflammation and neurodegenerative diseases, making them prime targets for therapies (15). Expression of *KPNA7* is highly restricted to oocytes and early embryogenesis where recessive mutations cause congenital skeletal abnormalities (16) and preimplantation embryo arrest (PREMBA) (17). Perhaps most significant to otosclerosis, Imp-αs are known to inhibit the nuclear import of parathyroid related protein (PTHrP), a major regulator of chondrogenesis (18). We explore how embryonically expressed genes such as *KPNA7* may cause adult-onset HL and garner insights into the pathobiology of otosclerosis and why the rest of the human skeleton is likely spared from abnormal bone deposition and disease.

## Results

### Clinical recruitment, pedigree structure and classification of hearing loss

The proband (PID III-1) was diagnosed at 42 with severe HL due to otosclerosis in both ears, which began in teenage years. Pre-surgery audiogram revealed bilateral, conductive HL with borderline cochlear (sensorineural) loss. Hearing improved bilaterally after two successive stapedectomies (Fig 1). The diagnosis of otosclerosis was confirmed upon surgical visualization of stapes fixation. Based on the medical questionnaires, HL started in the teens for all but one sibling who noticed HL in adulthood, and hearing was restored in all siblings after stapedectomy surgery. The pedigree structure is consistent with both AD and autosomal recessive (AR) inheritance, but X-linked inheritance can be ruled out as otosclerosis is not more severe in males.

**Fig 1.**
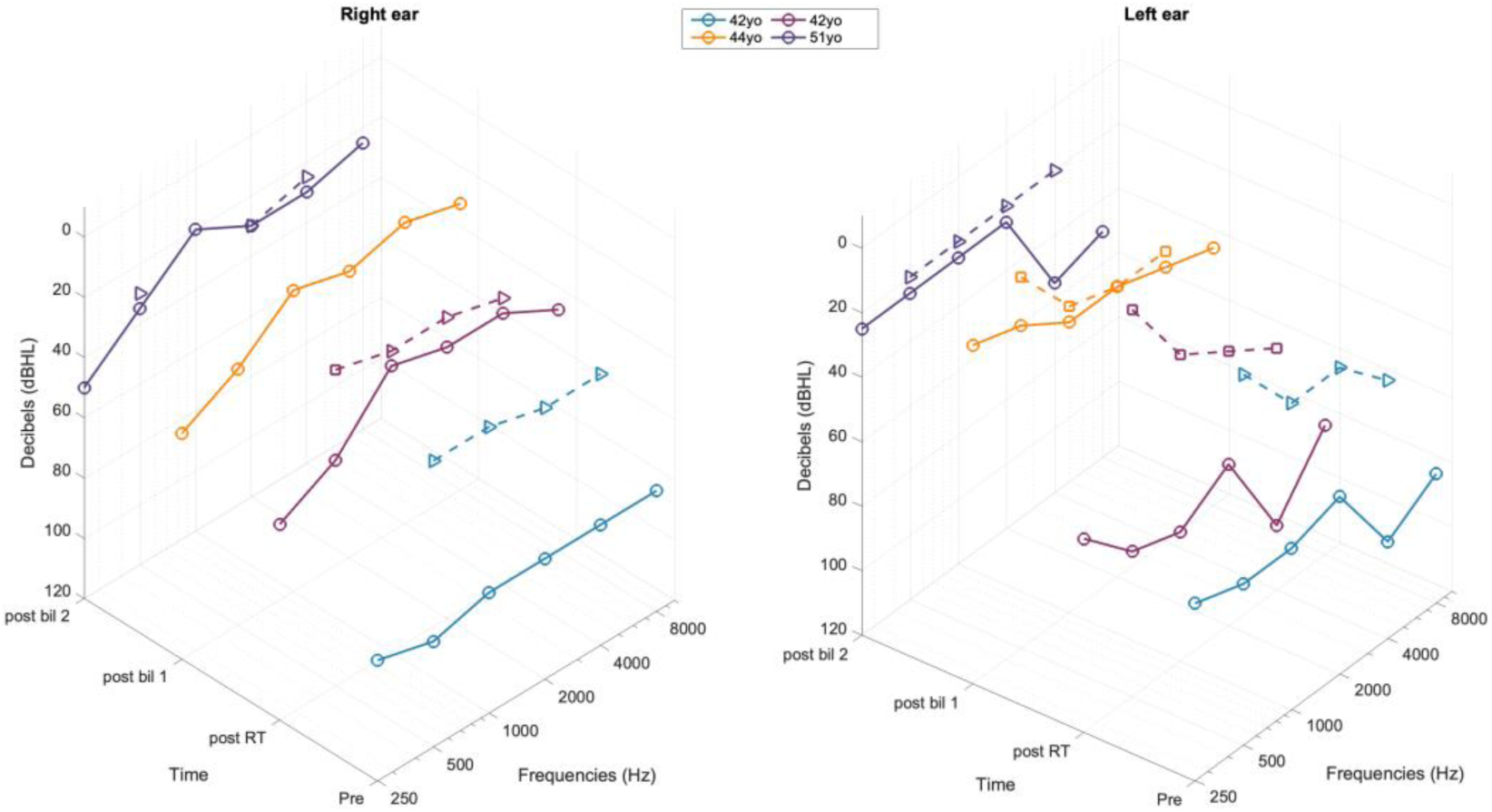
Pre- and post-stapedectomy audiograms reveal successful surgical treatment of bilateral otosclerosis in the proband (PID III-1). Pre-surgery audiogram revealed a conductive loss with borderline cochlear (sensorineural) HL. Post stapedectomy of the right ear improved hearing and the air-bone gap was mainly resolved but some HL remained, especially in the low frequencies. After stapedectomy of the left ear at age 44, hearing improved across all frequencies with only a mild sensorineural HL remaining in the mid to high frequencies. At age 51, the proband experienced mild hearing loss in the low and mid frequencies of the left ear but high frequencies showed moderate to severe loss at 4000 Hz and 8000 Hz, likely due to disease processes on the cochlear side of the round window. Pre = pre-stapedectomy audiogram, age 42; post RT = post-stapedectomy audiogram, right ear, age 42; post bil 1 = first post-bilateral stapedectomy audiogram, age 44; post bil 2 = second post-bilateral stapedectomy audiogram, age 51; ρ = unmasked bone conduction; □ = masked bone conduction. Generated by MathWorks. (2020). *MATLAB (Version R2020a)*[Computer software].

### Haplotype sharing on chromosome 7q and targeted gene sequencing

When testing for linkage to mapped *OTSC* loci and recapitulating disease-associated haplotypes, we would expect, under an AD model, to observe a single otosclerosis-associated haplotype that is shared exclusively among affected family members. Conversely, under an AR model, we would expect all affected to share the same maternal and paternal disease haplotypes. Although we have limited clinical data on the paternal side, under the AD model, we observed a shared paternal disease haplotype encompassing *OTSC2* (7q) (Fig 2). Conversely, no sharing was observed for *OTSC1* (15q), *OTSC3* (6p)*, OTSC4* (16q), *OTSC5* (3q), *OTSC7* (6q), *OTSC8* (9p) or the *COL1A1* (17q) and *NOG* (17q) genomic regions. Subsequent targeted gene sequencing of 11 positional candidate genes within the disease interval on 7q did not identify the otosclerosis gene. The proband also screened negative for the 15-base pair (bp) deletion in *FOXL1* (rs764026385; *OTSC11*) that we previously identified in a Newfoundland family, and for rare otosclerosis variants in *SERPINF1*.

**Fig 2.**
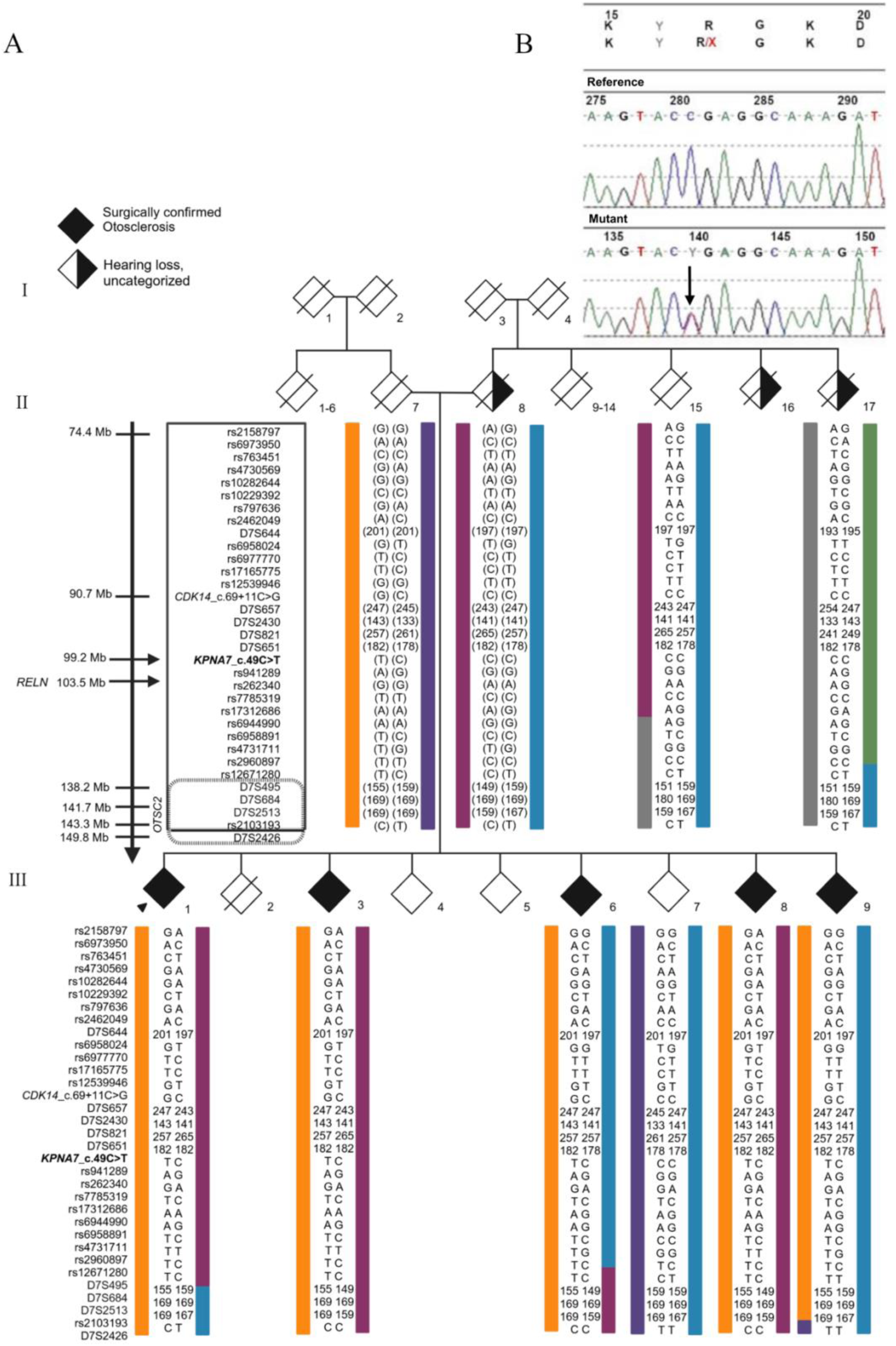
Pedigree and otosclerosis-associated haplotype of a multiplex family from NL with clinically confirmed otosclerosis. (A) Pedigree of NL family with AD otosclerosis co-segregating with the *KPNA7*, c.49C>T premature stop mutation and transmitted from the paternal side (orange haplotype). *KPNA7* maps to the *OTSC2* locus in the vicinity of *RELN.* (B) Sequencing electropherogram generated from Mutation Surveyor software showing the heterozygous nonsense mutation. We used diamonds instead of squares (males) and circles (females) to protect the identity of research participants.

### Genome wide analysis under AR linked regions

Although there was evidence for haplotype sharing on 7q, targeted gene sequencing did not identify the gene. Under an AR model, linkage simulation derived the theoretical maximum LOD score (LODmax=2.5) given the pedigree structure. SNP genotyping and multipoint linkage analysis yielded LODobs=2.5 at chr17q25.1-q25.3, spanning a region of 5.9 Mb. Exome sequencing of the 141 positional candidate genes revealed zero homozygous variants and 49 heterozygous variants. Only two genes, *TEN1* and *EVPL,* had two or more variants consistent with AR inheritance however, these were filtered out due to high population frequencies and/or benign functional predictions (19).

### Genotype wide analysis under AD linked regions

Assuming AD inheritance, the LODmax (1.73) was not obtained but suggestive LOD (LOD > 1) scores were observed at five distinct genomic loci (7q, 10p, 10q, 16q, 17q). Several of these overlapped with *OTSC* disease intervals, specifically *OTSC2* (7q), *OTSC4* (16q), *NOG* ( 17q). We identified 542 positional candidate genes under the five suggestive loci. Exome sequencing yielded 301 variants and of these,153 variants were filtered out as they were not identified in all affected. We could remove 37 variants as they were also identified in “solved” *FOXL1* cases. Of the remaining variants, 69 had MAF >2%, reducing the variants of interest to 23 silent, 15 missense, one intronic and one nonsense. The 15 missense mutations were all predicted to be benign, and 16 (15 silent, one intronic) were predicted to have no effect on splicing. Only the nonsense mutation in *KPNA7* gene on 7q (*OTSC2*) remained after variant filtering.

### Sanger validation, cascade sequencing and *in silico* analyses

*KPNA7*, c.49 C>T (NM_001145715.3) is rare (0.015%, gnomAD, rs746784660) and has been reported in ClinVar as a VUS (ID: 652650). We found that *KPNA7*, c.49 C>T is absent in 187 HL probands and in controls. Cascade sequencing confirmed co-segregated with otosclerosis (Fig 2). *KPNA7*, c.49 C>T (NM_001145715.3) is located near the 5’ end of *KPNA7* and is predicted to cause premature stop mutation, p. R17X, subjecting the truncated RNA transcript to nonsense mediated decay and resulting in haploinsufficiency (Alamut Visual Plus, version 1.13, Sophia Genetics (2024)). Previously, recessive mutations in *KPNA7* cause congenital skeletal abnormalities (16) and preimplantation embryo arrest (PREMBA) (17). We conclude that *KPNA7*, c.49 C>T, p. R17X is pathogenic according to HL ACMG criteria PVS1 and PM2 (20, 21).

## Discussion

### Summary of findings

We identify *KPNA7,* the newest of the human Imp-α transport factors, as the first of the nuclear transport system of proteins to cause AD otosclerosis. Surgical reports in a white multiplex family of Northern European extraction confirmed the diagnosis and affected siblings reported HL as young adults. Using comprehensive genetic and genomics analyses, we identified a disease-associated haplotype on chromosome 7q22.1 overlapping the *OTSC2* locus (22). The absence of paternal DNA was overcome by recruiting maternal siblings, confirming paternal transmission of a premature stop mutation in *KPNA7* [(NM_001145715.3), c.49 C>T, p. R17X]. *KPNA7* is located within the *OTSC2* locus and resides in the vicinity of *RELN* (OMIM 605727), a gene whose intronic variants have been validated in GWAS otosclerosis case studies (4, 23). Although not normally expressed in adult tissues, KPNA7 is reactivated in cancer cell lines, with the highest expression detected in pancreatic cell lines harbouring an amplification of the 7q21-22 genomic locus where KPNA7 resides (24).

### Otosclerosis may be the failure of two or more parallel protective mechanisms

Bloch and colleagues have eloquently modeled how age-dependent microdamage accumulates in the human perilabyrinthine bone where bone remodeling is essentially absent (13, 25–29). Clusters of dead osteocytes (cellular voids) lose connection with each other as the perilacunar matrix breaks down, rendering OPG and other nutrients beyond the reach of viable osteocytes trapped within these voids (26). In their model of aging, quiescent cells within the globuli interossei may be able to break free of suppressive actions of OPG and go on to complete the process of endochondral ossification. However, the accumulation of voids alone is unlikely to be causative as clinical otosclerosis is not prevalent (0.30–0.38% in Europeans) and histologic otosclerosis has been observed in under 3% of temporal bone autopsy specimens (30). Tissue from patients with labyrinthitis ossificans, a rare reaction to infection where the membranous labyrinth turns to bone, essentially halting OPG production, show the persistence of cartilage remnants and the absence of bone remodeling, suggesting parallel mechanisms, in addition to OPG, protect the otic capsule from remodeling (27). Our exploration into the function of KPNA7 in embryonic cells suggests that KPNA7 represents a parallel protective mechanism.

### The structure of Imp-αs and their many roles in nuclear transport

Cells require an active transport machinery to shuttle RNA, protein and other macromolecules to their correct subcellular localizations to maintain homeostasis and carry out normal functions. For proteins, the most utilized nuclear import pathway is mediated by Imp-αs (31). *KPNA7* encodes the newest of the seven-member imp-α karyopherins, a group of highly conserved proteins that share a common structure consisting of a body of ten helical Armadillo (ARM) repeats, a short C-terminal region of acidic amino acids and an N-terminal, Importin-β (Imp-β) binding (IBB) domain. Most proteins targeted for transport into the nucleus contain a nuclear localization signal (NLS) motif, a lysine-rich stretch of basic amino acids containing one (monopartite) or two (bipartite) basic regions separated by a linker region in their amino acid sequence (31). The first NLS motif to be recognized and best characterized is the classical NLS (cNLS). Imp-αs function as adaptors that recognize and bind to cNLS in their cargo proteins. Proteins with non-classical NLS (ncNLS) motifs can also be bound and transported directly by Imp-βs without the need for Imp-α adaptors.

Except for KPNA7, Imp-αs are maintained in a closed state (autoinhibited) in the cytoplasm with the highly flexible IBB domain folding back onto itself blocking the NLS binding groove from binding NLS-containing cargo (32, 33). In cNLS transport, the IBB domain binds Imp-β, exposing the NLS binding groove for specific cargo protein binding. The cargo-Imp-α-β tricomplex, once formed, is rapidly transported to the nucleus via Imp-β interactions with select nucleoporins lining the central channel of the nuclear pore complex. In ncNLS transport, Imp-β binds and transports NLS-containing cargo in the absence of Imp-α. Inside the nucleus, Imp-β binds RanGTP and releases Imp-α and cargo protein and then Imp-α and Imp-β are recycled back to the cytoplasm by export receptors.

Imp-αs can also act as negative regulators for the nuclear import of certain proteins by competing with Imp-β for NLS binding to cargo, or by forming a transport-incompetent complex in the cytoplasm, preventing cargo from entering the nucleus. For example, in the presence of Imp-α, TRF1 forms a complex with Imp-α-β but this complex remains in the cytoplasm. In contrast, Imp-α competes with the binding of imp-β to Snail zinc finger domain, resulting in ineffective nuclear accumulation of Snail, leading to a decrease in its cellular protein level through subsequent degradation by the protease system, with implications for the prevention of tumor cell invasion by inhibiting Snail localization (34). Perhaps most significant to otosclerosis, Imp-αs are known inhibitors of PTHrP, a multifunctional cytokine and a major regulator of chondrogenesis in the human skeleton sharing structural similarities with, and the same receptor (PTH1R) as parathyroid hormone (PTH) (35). Interestingly, PTH1R mRNA expression in otosclerotic stapes led Grayeli and colleagues to hypothesize that abnormal cellular response to PTH played a role in abnormal remodeling in otosclerosis (36).

### Characteristics of KPNA7, the newest nuclear import factor

KPNA7 is the most recent and divergent of the seven human Imp-αs and an intriguing *OTSC* gene as it is not normally expressed in adult tissues. KPNA7 is recognised as a maternal factor essential for embryogenesis and fertility (37) and plays a critical role in protein transport in oocytes and early embryos and is critical to early embryonic cleavage events and zygotic genome activation (38). *KPNA7* is known to have many cargo proteins, up to 377 have been reported (24). KPNA7 has a unique ability among the Imp-αs to maintain an open state and has the strongest IBB domain capacity for Imp-β, likely critical to producing pre-formed Imp-α-β heterodimers to increase the transport rate of cargo proteins in the early stages of embryogenesis (38). KPNA7 is the most abundant Imp-α in germinal vesicle and metaphase II-stage oocytes (39) and is nearly absent from the eight-cell embryo onward (38), being rapidly degraded during zygotic genome activation and barely detectable in morula- and blastocyst-stage embryos (40). Recessive *KPNA7* mutations result in low protein expression levels, interfering with the nuclear import of RSL1D1 (also known as cellular senescence-inhibited gene protein: CSIG) (17). RSL1D1 negatively regulates PTEN via translational suppression causing increased cell proliferation, is significantly elevated in the tumors of colorectal cancer patients predicting poorer survival outcomes and is a potential new target for cancer therapies (41). Imp-α has been shown to inhibit ncNLS transport of certain proteins including PTHrP (42). As the N-terminal domain of Imp-β binds PTHrP at HEAT repeats 2-11, but also binds Imp-α (IBB domain) at HEAT repeats 7-19, this partial overlap of binding sites may explain ncNLS transport inhibition of PTHrP (43).

### Nuclear import of PTHrP drives developmental pathways in a context-specific manner

The otic capsule forms through a series of molecular and cellular signaling processes where mesenchymal progenitor cells undergo condensation and differentiation into chondrocytes which subsequently proliferate and hypertrophy, followed by mineralization of the extracellular matrix and apoptosis of the chondrocytes (44) (Fig 3). PTHrP is necessary for endochondral ossification, regulating chondrocyte maturation, proliferation and differentiation (44). In fact, PTHrP is essential to development. PTHrP null mice (Pthrp −/−) die early in the postnatal period and display severe chondrodysplasia with reduced endochondral development and excessive mineralization (45). Experiments testing the effect of mechanical strain on chondrocytes showed that PTHrP expression increased during the proliferation and matrix forming stages under conditions of cyclical strain due to Indian Hedgehog (Ihh) signaling in chondrocytes (46). The Ihh signaling pathway also works with other signaling molecules and pathways to promote chondrocyte proliferation and inhibit hypertrophy and forms a negative feedback loop with PTHrP pivotal in regulating cartilage development (47). Upon activation of Ihh expression from an external trigger such as mechanical stress, Ihh binds to the transmembrane protein Patched (Ptc), releasing inhibition of Smoothened (Smo) at the cell surface, which then activates the expression of the downstream signaling molecule Gli. Gli enters the nucleus and regulates the expression of downstream signaling factors SOX9, RUNX2 and PTHrP (48) (Fig 3).

**Fig 3.**
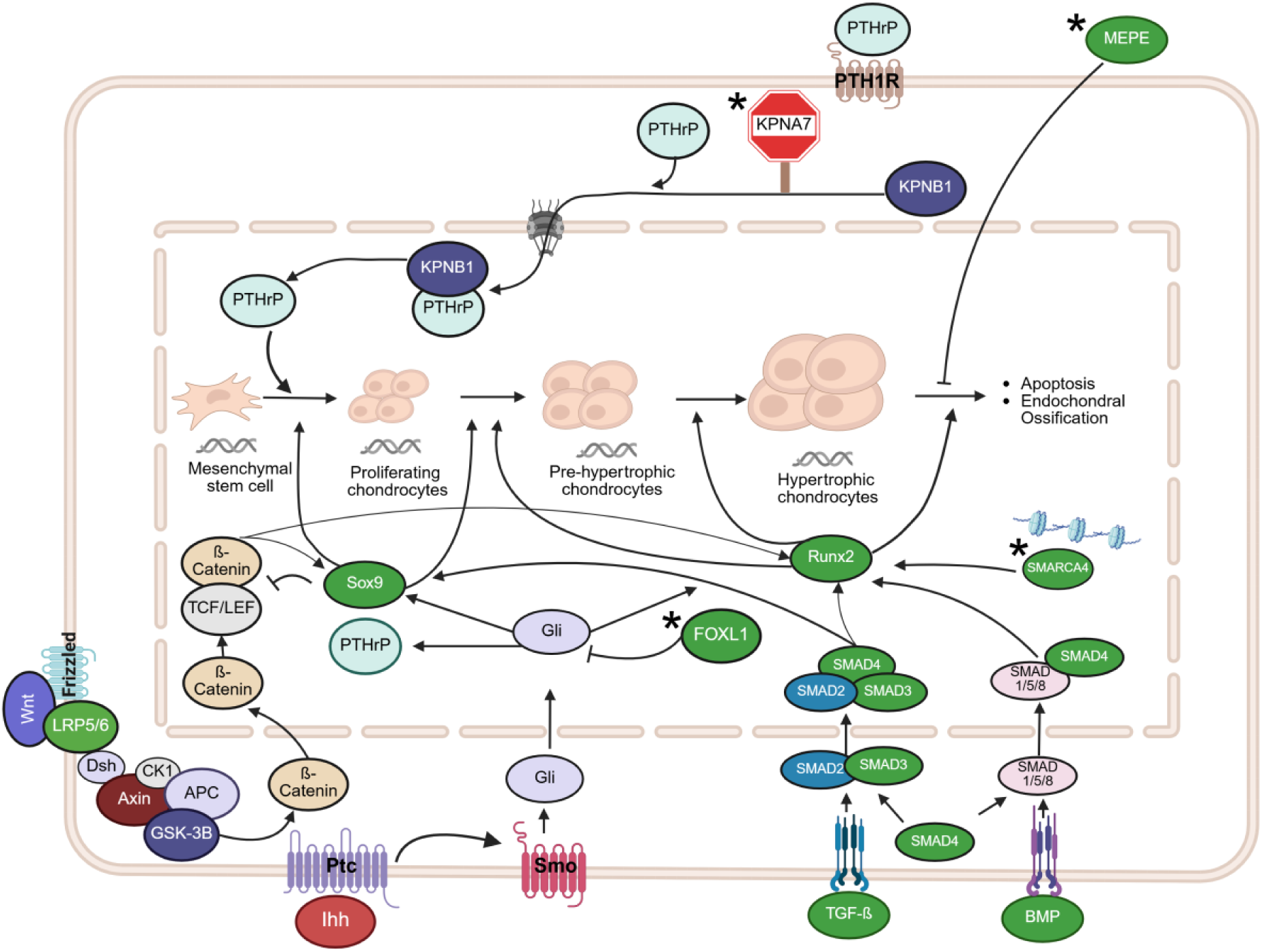
New mechanistic model for otosclerosis due to pathogenic mutations in *KPNA7* and other *OTSC* genes within the aging otic capsule. Otosclerosis genes (*) and otosclerosis susceptibility genes (green circles) are involved in endochondral ossification signaling pathways. We hypothesize that KPNA7 haploinsufficiency allows the nuclear import of PTHrP into the quiescent cells of globuli interossei trapped within cellular voids of the aging otic capsule, re-activating endochondral ossification pathways in these embryonic tissues. Created in BioRender. BENTEAU, T. (2025) https://BioRender.com/mfhqhzv.

### How KPNA7 haploinsufficiency may cause otosclerosis

As PTHrP nuclear import is integral to its function in cells, strategies to block PTHrP nuclear import could have important effects on target cell function (18). Endochondral ossification is initiated by nuclear import of PTHrP and quiescent chondrocytes are known to express PTHrP (49). Upon entering the nucleus by ncNLS transport chondrocyte regulation is achieved through the Ihh-PTHrP pathway (44), as well as the WNT/ β-catenin pathway (50) via negative feedback loops. PTHrP is both fast acting and fast to be removed (51). In a scenario where the KPNA7-Imp-β complex is pre-formed, potentially KPNA7 would be able to prevent binding of PTHrP by binding a different cargo or outcompete PTHrP and enter the nucleus in a cargo-free state, thereby inhibiting nuclear transport of PTHrP, as suggested by Oostdyk and colleagues for other cargo proteins (38). While KPNA7 has a time-limited role in the nuclear import of factors required for maternal-to-zygotic transition and early embryogenesis, it is logical to think that KPNA7 also inhibits endochondral ossification by preventing nuclear import of PTHrP in the unique and peculiar case of the embryonic cells of the globuli interossei.

Although not normally expressed in adult tissues, KPNA7 is reactivated in cancer cell lines promoting carcinogenesis by increasing the rate of import of cell cycle factors due to genomic amplification on 7q21-22. Unlike amplification in pancreatic cancer, the effect of a heterozygous KPNA7 stop mutation identified in otosclerosis patients is predicted to cause its phenotypic effects via haploinsufficiency. Therefore, it is unlikely that otosclerosis is due to increased import of KPNA7 cargo proteins. The quiescent chondrocytes of the globuli interossei, unique to the otic capsule, would provide a likely substrate for KPNA7 (10, 11). While a mutation in an Imp-α exclusively expressed in early development would not be expected to impact the rest of the human skeleton, we hypothesize that KPNA7 plays a key role in preventing PTHrP nuclear import in quiescent chondrocytes. If this is indeed the case, KPNA7 haploinsufficiency may activate chondrogenesis leading to the deposition of spongy, disorganized bone characteristic of osteosclerotic specimens, and specifically, the mineralization stage of the extracellular matrix in hypertrophic chondrocytes. If this speculation is true, otosclerosis is not the result of pathologic bone resorption and deposition, but one of re-activation of chondrogenesis, providing an alternative mechanism for its pathogenesis.

### OTSC genes and their roles in endochondral ossification

A closer look at the *OTSC* genes *MEPE*, *FOXL1* and *SMARCA4* reveal they are involved in endochondral ossification signaling pathways (Fig 3). In addition, the *OTSC* gene *SERPINF1* encodes for PEDF, which is involved in many biological processes including bone formation, binding to extracellular matrix proteins including collagen and glycosaminoglycan, and is involved in the mineralization of bone matrix (52). Rämö and colleagues identified otosclerosis susceptibility genes BMP-2,-3,-4,-7 (involved in bone development) and SOX9 and RUNX2 (4). MEPE normally serves as a decoy receptor for pre-osteoclasts inhibiting osteoclast maturation and its ASARM motif, upon proteolytic cleavage by β-cathepsin, inhibits mineralization by binding to hydroxyapatite crystals (53). In zebrafish, foxl1 (*OTSC11*) regulates the expression of collagen genes such as col1a1 and col11a2, and results in a delay in jawbone mineralization (54). SMARCA4 (also known as BRG1) plays a role in ossicle formation during embryogenesis and may be important for regulation of osteoblast differentiation and maintenance of postnatal bone homeostasis in the otic capsule (55). WNT/ β-catenin is also involved in proliferation and differentiation through a negative feedback loop with PTHrP (50).

### Re-activation of endochondral ossification pathways in the globuli interossei, a new mechanistic model for otosclerosis

Otosclerosis is likely due to germline susceptibility factors interacting with environmental triggers (measles infection, pregnancy, etc.) to generate the pathological phenotype (2). Perhaps deficient KPNA7 causes the reactivation of chondrogenesis and the resumption of endochondral ossification after loss of repression by OPG in aging otic capsules. From this perspective, the pathologic spongy bone deposition seen in otosclerosis may be the result of dysregulation caused by mutations in *OTSC* genes regulating endochondral ossification. Endochondral ossification involves multiple signaling pathways including the Ihh pathway, which regulates chondrocyte maturation and bone formation, the PTHrP pathways and bone morphogenetic proteins (BMPs). Ihh regulates chondrocyte maturation and bone formation, and bone morphogenetic proteins (BMPS) induce chondrocyte differentiation via regulating the expression of SOX9 and stimulating endochondral ossification via transcriptional regulation of RUNX2.

### Could dysregulation of PTHrP due to KPNA7 cause otosclerosis in women

In a literature review on otosclerosis and pregnancy, Fabbris and colleagues found that the only significant correlation was between pregnancy and disease onset, noting that cases with hearing impairment increased with number of pregnancies (56). We know that PTHrP levels are higher in pregnant women, and are significantly higher in lactating women, being detected in breast milk at levels exceeding 10,000 times those found in the blood of hypercalcemia of malignancy patients or normal controls (57). Otosclerosis during pregnancy may be the result of increased bone remodeling (RANK/RANKL/OPG) (57–59) or due to susceptibility variants in KPNA7 or other Imp-αs involved in regulating the actions of PTHrP.

### Overlapping therapeutic targets in skeletal disorders and cancer

*OTSC* genes *FOXL1*, *SMARCA4* and *KPNA7* are increasingly recognised for their role in carcinogenesis (60–64). By extension, therapeutic targets for cancer, otosclerosis and other skeletal disorders increasingly overlap because the development, proliferation and migration of cancer cells mimic critical pathways of embryonic development. This includes deregulation in nucleocytoplasmic transport. For example, expression of KPNA7 in adult tissues is almost non-existent, except in pancreatic cancer cell lines where KPNA7 causes a significant decrease in cell growth due to G1 arrest, accompanied by an increased expression of p21, a key regulator of the cell cycle (65). As we have seen, low KPNA7 levels causes PREMBA through dysregulated import of RSL1D1, which in turn, negatively regulates PTEN and is a potential new target for colorectal cancer (41). Imp-αs also facilitate the nuclear import of Smad proteins in the TGF-β/Smad3 pathway and show promise as therapeutic targets in rotator cuff injuries (66). There is an epidemic of colorectal cancer in young people (67) and of adult-onset skeletal disorders causing a public health issue in the global North. An investment in therapeutic options for cancer and for the repair of cartilage for aging knees and hips may benefit patients with otosclerosis by providing target drugs for off-label clinical trials.

### Limitations of this study

Although *KPNA7* maps to the *OTSC2* locus (22), we did not have access to the family used to map *OTSC2*, therefore we cannot confirm or deny that *KPNA7* is *OTSC2*. Association between otosclerosis and *RELN* may be due to the close physical proximity of *RELN* to disease-variants in *KPNA7*. We assume equal expression of maternal and paternal *KPNA7* alleles in the globuli interossei however we do not have patient-derived disease tissue to examine the functional consequences of *KPNA7*, c.49C>T, p.R17X. Otosclerosis due to KPNA7 haploinsufficiency may be related to the dysregulated import of RSL1D1 or there may be alternative mechanistic explanations for the pathogenesis of KPNA7 as the specific binding affinities and cargo preferences of KPNA7, being the most recent member of the Imp-αs, have not been fully explored. Evidence from this study suggests that reactivation of endochondral ossification, not the de-repression of bone remodeling, underlies otosclerosis due to *KPNA7* mutation. Future research is warranted along this direction to describe the modulators of KPNA7 function and to design potential therapies to modulate the PTHrP-Imp-α interactions for therapeutic purposes.

### Future directions

We used strict clinical criteria and a combination of genetic and genomic analyses to successfully identify a novel *OTSC* gene, *KPNA7*, the second *OTSC* gene identified in the white settler population of the island of Newfoundland, Canada. KPNA7 is the first member of the nucleocytoplasmic trafficking system to cause otosclerosis. We propose that normally, *KPNA7* plays a critical role of inhibiting the nuclear import of PTHrP in the globuli interossei and that KPNA7 haploinsufficiency results in reactivation of endochondral ossification, providing an additional or alternative paradigm to the central hypothesis that dysregulation of bone remodeling repression causes otosclerosis. As the molecular signaling pathways involved in embryogenesis also play a role in the fate of neoplastic cells, investigating the shared cellular and molecular signaling pathways and molecules that cause other monogenic skeletal disorders and carcinomas may provide unique insights into novel therapeutic targets for common maladies including otosclerosis, osteoarthritis and cancer. Difficulties with accessing the complex, fluid-filled cavity of the inner ear within the dense temporal bone, currently hampering diagnosis and targeted drug delivery, may be behind us with the advent of microneedle technologies for safe sampling of inner ear fluids and local treatments, as microlitres of perilymph fluid can be extracted this way (68). The class of molecules that do not encode for proteins but regulate gene expression, non-coding RNA, are revealing an increasing role in cellular process, including differentiation and maturation, and their dysregulation can cause cancer. Perhaps renewed interest in solving the *OTSC* loci should include looking for non-coding RNA targets.

## Materials and Methods

### Ethics Statement

Prior approval to study live research participants with HL and their blood relatives was granted by the regional ethics authority (Hearing Loss Project #01.186, Human Research Ethics Board, St John’s, NL, Canada). Informed consent from research participants was by written consent.

### Clinical recruitment, pedigree structure and classification of HL

Research participants underwent audiological and medical examinations, permitted access to their medical records, and completed a medical questionnaire (19). HL was classified based on the pure-tone threshold averages of 0.5, 1.0 and 2.0 kHz, as defined by the American Speech and Hearing Association (AHSA). Family members were assessed by our clinical team to update audiograms and confirm middle ear status. A difference of >10 dB HL between air and bone conduction sensitivity represented a significant conductive component associated with impaired sound transfer through the middle ear. Conservative clinical criteria were used to assign otosclerosis affection status: affected were blood relatives with surgically confirmed otosclerosis at any age; unaffected were blood relatives ≥ 55 years of age with normal bilateral hearing thresholds. Of nine siblings in generation III, five have otosclerosis (Fig 2). The father (PID II-7) reportedly had normal hearing but was “very sensitive to noise” and the mother (PID II-8) had HL (uncategorized) along with two of her nine siblings (PID II-16, PID II-17) who are reported to have age-related HL. X-linked inheritance could be ruled out as otosclerosis is not more severe in males; however, without further clinical information, the inheritance pattern is consistent with both AD and AR inheritance.

### Test for linkage to mapped OTSC loci-haplotype sharing and targeted sequencing

Genomic DNA was extracted from peripheral blood (69). Informative microsatellite markers (4-10 per locus) spanning seven *OTSC* loci and two otosclerosis-suceptibility genes including *OTSC1* (15q), *OTSC2* (7q)*, OTSC3* (6p), *OTSC4* (16q), *OTSC5* (3q), *OTSC7* (6q), *OTSC8* (9p) and genes *COL1A1* (17q) and *NOG* (17q) were genotyped and shared alleles noted among affected siblings. Primers were labelled (6-FAM), amplified using touchdown PCR, size fractionated (ABI PRISM model 3130xl) and analyzed with Gene Mapper software (v4.0). Haplotypes were recapitulated according to “least recombination rules” and paternal haplotypes were inferred due to absence of DNA. Full gene sequencing (Sanger) was performed on three affected (PIDs III-1, III-3, III-9) and one unaffected (PID III-7) for select annotated genes [March 2006 assembly (NCBI build 36.1)] within *OTSC* loci with disease-associated haplotypes. Genes were selected for sequencing if they had a functional role in bone remodeling or immune response. Primers were designed to amplify the longest isoform, including all intron/exon boundaries and UTRs and purified PCR products were bidirectionally sequenced (Big Dye Terminator V3.1 kit; ABI PRISM 3130xl DNA Analyzer). We used Mutation Surveyor software (version 4.07, SoftGenetics LLC State College, PA 16803) to select quality reads and analyze DNA sequences. Rare sequencing variants (<2 %) were subjected to *in silico* tools (SIFT, PolyPhen, Human Splicing Finder (HSF), MaxEntScan, NNSLICE, GeneSplicer, Known constitutive signals) to predict pathogenicity. Variants were filtered out if they were absent in affected or present in unaffected relatives. The proband was also screened for the 15-bp coding deletion in *FOXL1* (rs764026385; *OTSC11*) identified in an NL family (70) and for the rare otosclerosis variants in *SERPINF1* (71). All remaining variants were subjected to cascade sequencing to verify co-segregation with otosclerosis in the family.

### Genome wide SNP genotyping and multipoint linkage analysis

As a targeted genetic analysis did not identify the otosclerosis gene and the inheritance pattern is not clear, genome wide SNP genotyping and multipoint linkage analysis were performed under both AR and AD models with Merlin (version 1.1.2) (72, 73), assuming complete (100%) penetrance. LOD scores were calculated at recombination fractions of 0.000 to 0.5000. We used the 610K Illumina SNP array (Genome Centre, McGill University, QC, Canada) on samples from five affected siblings (PIDs III-1, III-3, III-6, III-8, III-9), an unaffected sibling (PID III-7), a maternal sibling with HL (PID II-17) and a maternal sibling with normal hearing (PID II-15). As well, nine population control samples with normal hearing were used to estimate minor allele frequencies of unavailable family members. Genotypes were analyzed at The Centre for Applied Genomics (TCAG, University of Toronto, ON, Canada) and exported from GenomeStudio software (v2010.3).

### Sequencing, variant filtering and cascade sequencing under linked regions

Exome sequencing was carried out on four affected siblings (PIDs III-1, III-3, III-6, and III-9) and two older controls (55, 60 yrs old) with normal hearing thresholds. Library preparation was done with TrueSeq Prep Kit and samples run on the Illumina Hiseq 2000, generating 50-150 million 100-bp paired end reads. Reads >32-bp long were aligned to the 1000 genome reference using Burrows-Wheeler Aligner (BWA) and merged with Picard software (Broad Institute). Where multiple base mismatches and false positive variant calls were recorded, insertions and deletions were realigned using GATK software (73, 74). The percentage of aligned region coverage was detected using the Genome Centre’s in-house database. The regions were identified as high coverage (>400X), low coverage (<50X), low mean mapq MQ (<20X) and no data. Rare variants were considered at a higher frequency (MAF<2%) to account for potential founder effects. Rare variants, absent in one or more affected, present in one or more controls or also identified in solved otosclerosis patients (e.g. *FOXL1*) were filtered out. Variants with a minimum of 20X coverage were analysed with *in silico* tools including samtools mpileup algorithm (75), SnpSift (76), SnpEff (76), SIFT, PolyPhen-2, PANTHER and ClustalW. In addition, the functional consequence of *KPNA7*, c.49 C>T, p. R17X was analysed using Alamut Visual Plus by Sophia Genetics, version 1.13 (2024). At this stage, allele frequencies of genetic variants were checked in two research cohorts, the NL Osteoarthritis Study (NFOAS) consisting of 1,000 total joint (knee and/or hip) replacement patients and the NL Colorectal Cancer Registry, where samples were genotyped by Illumina microarray platforms and then imputed with 1000 genome project data as reference panels. All remaining variants underwent cascade screening and were tested against 38 otosclerosis probands (from ON Canada; Western University ethics #103,679), 149 uncategorized HL probands (from NL) and 727 population controls.

## Web Resources

ASHA (Amer Speech-Language Hearing Assoc), https://www.asha.org/practice-portal/clinical-topics/hearing-loss/

Burrows-Wheeler Aligner BWA, http://bio-bwa.sourceforge.net/

ClinVar, https://www.ncbi.nlm.nih.gov/clinvar/

ClustalW, https://www.genome.jp/tools-bin/clustalw

dbSNP, https://www.ncbi.nlm.nih.gov/snp/

gnomAD, https://gnomad.broadinstitute.org/

Hereditary Hearing Loss homepage, http://hereditaryhearingloss.org/

Human Splicing Finder, www.umd.be/HSF3/

1000 genomes, https://www.internationalgenome.org/home

Online Mendelian Inheritance in Man, http://omim.org

Primer3, https://bioinfo.ut.ee/primer3-0.4.0/

PANTHER, http://www.pantherdb.org/

PolyPhen-2, http://genetics.bwh.harvard.edu/pph2/

RefSeq, https://www.ncbi.nlm.nih.gov/refseq/

SIFT, https://sift.bii.a-star.edu.sg/

SNP database, http://www.ncbi.nlm.nih.gov/projects/SNP/

The MathWorks, Inc. https://www.mathworks.com

Varsome, http://varsome.com

UCSC Genome Browser, https://genome.ucsc.edu/

## Acknowledgements

This study was funded by the Canadian Institutes of Health Research (#222294), Canadian Foundation for Innovation (#9384, #13120), and Genome Canada/Genome Atlantic (AMGGI) to T.L.Y. Support was also provided by Memorial University, Town of Grand Falls-Windsor (Excite Corporation) and the Government of Newfoundland and Labrador. One of the first authors, N.A., is a recipient of a CIHR Fellowship and this paper includes contributions from their PhD research.

## References

1. Sönmez S, Orhan KS, Baumgartner W-D, Zarowski A, Özgirgin NO, Barbara M, et al., editors. the Fifth International Symposium on Otosclerosis and Stapes Surgery. Symposium on Otosclerosis and Stapes Surgery J Int Adv Otol; 2025.

2. Capobianco S, Lazzerini F, Bruschini L, Fiacchini G, Forli F. Exploring the genetic landscape of otosclerosis: current understanding and future perspectives. Acta Otorhinolaryngol Ital. 2025;45(Suppl. 1):S2–s17.

3. Mudry A. Adam Politzer (1835-1920) and the description of otosclerosis. Otol Neurotol. 2006;27(2):276–81.

4. Rämö JT, Kiiskinen T, Seist R, Krebs K, Kanai M, Karjalainen J, et al. Genome-wide screen of otosclerosis in population biobanks: 27 loci and shared associations with skeletal structure. Nat Commun. 2023;14(1):157.

5. Huybrechts Y, Mortier G, Boudin E, Van Hul W. WNT Signaling and Bone: Lessons From Skeletal Dysplasias and Disorders. Front Endocrinol (Lausanne). 2020;11:165.

6. Sanverdi SE, Ozgen B, Dolgun A, Sarac S. Incomplete endochondral ossification of the otic capsule, a variation in children: evaluation of its prevalence and extent in children with and without sensorineural hearing loss. AJNR Am J Neuroradiol. 2015;36(1):171–5.

7. Frisch T, Sørensen MS, Overgaard S, Lind M, Bretlau P. Volume-Referent Bone Turnover Estimated From the Interlabel Area Fraction After Sequential Labeling. Bone. 1998;22(6):677–82.

8. Moser T, Veillon F, Sick H, Riehm S. The hypodense focus in the petrous apex: a potential pitfall on multidetector CT imaging of the temporal bone. AJNR Am J Neuroradiol. 2008;29(1):35–9.

9. Hawke M, Jahn AF. Bone formation in the normal human otic capsule. Arch Otolaryngol. 1975;101(8):462–4.

10. Richard C, Doherty JK, Fayad JN, Cordero A, Linthicum FH, Jr. Identification of target proteins involved in cochlear otosclerosis. Otol Neurotol. 2015;36(5):923–31.

11. Wang PC, Merchant SN, McKenna MJ, Glynn RJ, Nadol JB, Jr. Does otosclerosis occur only in the temporal bone? Am J Otol. 1999;20(2):162–5.

12. Zehnder AF, Kristiansen AG, Adams JC, Merchant SN, McKenna MJ. Osteoprotegerin in the inner ear may inhibit bone remodeling in the otic capsule. Laryngoscope. 2005;115(1):172–7.

13. Hansen LJ, Bloch SL, Sørensen MS. Cellular voids in the pathogenesis of otosclerosis. Acta Otolaryngol. 2023;143(3):250–3.

14. Jáuregui EJ, Akil O, Acevedo C, Hall-Glenn F, Tsai BS, Bale HA, et al. Parallel mechanisms suppress cochlear bone remodeling to protect hearing. Bone. 2016;89:7–15.

15. Mallik S, Poch D, Burick S, Schlieker C. Protein folding and quality control during nuclear transport. Curr Opin Cell Biol. 2024;90:102407.

16. Paciorkowski AR, Weisenberg J, Kelley JB, Spencer A, Tuttle E, Ghoneim D, et al. Autosomal recessive mutations in nuclear transport factor KPNA7 are associated with infantile spasms and cerebellar malformation. Eur J Hum Genet. 2014;22(5):587–93.

17. Wang W, Miyamoto Y, Chen B, Shi J, Diao F, Zheng W, et al. Karyopherin α deficiency contributes to human preimplantation embryo arrest. J Clin Invest. 2023;133(2).

18. Martin TJ. Parathyroid Hormone-Related Protein, Its Regulation of Cartilage and Bone Development, and Role in Treating Bone Diseases. Physiol Rev. 2016;96(3):831–71.

19. Abdelfatah N. The genetic aetiology of otosclerosis in the population of Newfoundland and Labrador: Memorial University of Newfoundland; 2014.

20. Oza AM, DiStefano MT, Hemphill SE, Cushman BJ, Grant AR, Siegert RK, et al. Expert specification of the ACMG/AMP variant interpretation guidelines for genetic hearing loss. Hum Mutat. 2018;39(11):1593–613.

21. Richards S, Aziz N, Bale S, Bick D, Das S, Gastier-Foster J, et al. Standards and guidelines for the interpretation of sequence variants: a joint consensus recommendation of the American College of Medical Genetics and Genomics and the Association for Molecular Pathology. Genet Med. 2015;17(5):405–24.

22. Van Den Bogaert K, Govaerts PJ, Schatteman I, Brown MR, Caethoven G, Offeciers FE, et al. A second gene for otosclerosis, OTSC2, maps to chromosome 7q34-36. Am J Hum Genet. 2001;68(2):495–500.

23. Schrauwen I, Ealy M, Huentelman MJ, Thys M, Homer N, Vanderstraeten K, et al. A genome-wide analysis identifies genetic variants in the RELN gene associated with otosclerosis. Am J Hum Genet. 2009;84(3):328–38.

24. Vuorinen E. Nuclear import protein KPNA7 and its cargos: Diverse roles in the regulation of cancer cell growth, mitosis and nuclear morphology. 2018.

25. Bloch SL. On the biology of the bony otic capsule and the pathogenesis of otosclerosis. Dan Med J. 2012;59(10):B4524.

26. Bloch SL, Sørensen MS. The role of connectivity and stochastic osteocyte behavior in the distribution of perilabyrinthine bone degeneration. A Monte Carlo based simulation study. Hear Res. 2016;335:1–8.

27. Frisch T, Bloch SL, Sørensen MS. Prevalence, size and distribution of microdamage in the human otic capsule. Acta Oto-Laryngologica. 2015;135(8):771–5.

28. Hansen LJ, Bloch SL, Sørensen MS. Identification of cellular voids in the human otic capsule. Journal of the Association for Research in Otolaryngology. 2021;22(5):591–9.

29. Hansen LJ, Bloch SL, Frisch T, Sørensen MS. Microcrack surface density in the human otic capsule: An unbiased stereological quantification. The Anatomical Record. 2021;304(5):961–7.

30. Declau F, van den Bogaert K, Van de Heyning P, Offeciers E, Govaerts P, van Camp G. Phenotype-genotype correlations in otosclerosis: clinical features of OTSC2. Adv Otorhinolaryngol. 2007;65:114–8.

31. Lange A, Mills RE, Lange CJ, Stewart M, Devine SE, Corbett AH. Classical nuclear localization signals: definition, function, and interaction with importin α. Journal of Biological Chemistry. 2007;282(8):5101–5.

32. Kelley JB, Talley AM, Spencer A, Gioeli D, Paschal BM. Karyopherin alpha7 (KPNA7), a divergent member of the importin alpha family of nuclear import receptors. BMC Cell Biol. 2010;11:63.

33. Kobe B. Autoinhibition by an internal nuclear localization signal revealed by the crystal structure of mammalian importin alpha. Nat Struct Biol. 1999;6(4):388–97.

34. Sekimoto T, Yoneda Y. Intrinsic and extrinsic negative regulators of nuclear protein transport processes. Genes to Cells. 2012;17(7):525–35.

35. Lam MH, Thomas RJ, Martin TJ, Gillespie MT, Jans DA. Nuclear and nucleolar localization of parathyroid hormone-related protein. Immunol Cell Biol. 2000;78(4):395–402.

36. Grayeli AB, Sterkers O, Roulleau P, Elbaz P, Ferrary E, Silve C. Parathyroid hormone-parathyroid hormone-related peptide receptor expression and function in otosclerosis. Am J Physiol. 1999;277(6):E1005–12.

37. Pumroy RA, Cingolani G. Diversification of importin-α isoforms in cellular trafficking and disease states. Biochem J. 2015;466(1):13–28.

38. Oostdyk LT, McConnell MJ, Paschal BM. Characterization of the Importin-β binding domain in nuclear import receptor KPNA7. Biochem J. 2019;476(21):3413–34.

39. Sharif M, Detti L, Van den Veyver IB. Take your mother’s ferry: preimplantation embryo development requires maternal karyopherins for nuclear transport. J Clin Invest. 2023;133(2).

40. Tejomurtula J, Lee K-B, Tripurani SK, Smith GW, Yao J. Role of importin alpha8, a new member of the importin alpha family of nuclear transport proteins, in early embryonic development in cattle. Biology of reproduction. 2009;81(2):333–42.

41. Liu X, Chen J, Long X, Lan J, Liu X, Zhou M, et al. RSL1D1 promotes the progression of colorectal cancer through RAN-mediated autophagy suppression. Cell Death & Disease. 2022;13(1):43.

42. Miyamoto Y, Yamada K, Yoneda Y. Importin α: a key molecule in nuclear transport and non-transport functions. J Biochem. 2016;160(2):69–75.

43. Cingolani G, Bednenko J, Gillespie MT, Gerace L. Molecular basis for the recognition of a nonclassical nuclear localization signal by importin beta. Mol Cell. 2002;10(6):1345–53.

44. Chen H, Tan XN, Hu S, Liu RQ, Peng LH, Li YM, et al. Molecular Mechanisms of Chondrocyte Proliferation and Differentiation. Front Cell Dev Biol. 2021;9:664168.

45. Karaplis AC, Luz A, Glowacki J, Bronson RT, Tybulewicz V, Kronenberg HM, et al. Lethal skeletal dysplasia from targeted disruption of the parathyroid hormone-related peptide gene. Genes & development. 1994;8(3):277–89.

46. Tanaka N, Ohno S, Honda K, Tanimoto K, Doi T, Ohno-Nakahara M, et al. Cyclic mechanical strain regulates the PTHrP expression in cultured chondrocytes via activation of the Ca2+ channel. Journal of dental research. 2005;84(1):64–8.

47. Wu Q, Zhang Y, Chen Q. Indian hedgehog is an essential component of mechanotransduction complex to stimulate chondrocyte proliferation. J Biol Chem. 2001;276(38):35290–6.

48. Chen Y, Mehmood K, Chang Y-F, Tang Z, Li Y, Zhang H. The molecular mechanisms of glycosaminoglycan biosynthesis regulating chondrogenesis and endochondral ossification. Life Sciences. 2023;335:122243.

49. Hallett SA, Matsushita Y, Ono W, Sakagami N, Mizuhashi K, Tokavanich N, et al. Chondrocytes in the resting zone of the growth plate are maintained in a Wnt-inhibitory environment. Elife. 2021;10:e64513.

50. Tong W, Xu J, Qi Q, Chen H, Huang T, Chen C, et al. PTHrP buffers Wnt/β-catenin activity through a negative feedback loop to maintain articular cartilage homeostasis. bioRxiv. 2022:2022.11. 25.517940.

51. Schipani E, Lanske B, Hunzelman J, Luz A, Kovacs CS, Lee K, et al. Targeted expression of constitutively active receptors for parathyroid hormone and parathyroid hormone-related peptide delays endochondral bone formation and rescues mice that lack parathyroid hormone-related peptide. Proc Natl Acad Sci U S A. 1997;94(25):13689–94.

52. Al-Jallad H, Palomo T, Roughley P, Glorieux FH, McKee MD, Moffatt P, et al. The effect of SERPINF1 in-frame mutations in osteogenesis imperfecta type VI. Bone. 2015;76:115–20.

53. Schrauwen I, Valgaeren H, Tomas-Roca L, Sommen M, Altunoglu U, Wesdorp M, et al. Variants affecting diverse domains of MEPE are associated with two distinct bone disorders, a craniofacial bone defect and otosclerosis. Genet Med. 2019;21(5):1199–208.

54. Hawkey-Noble A, Pater JA, Kollipara R, Fitzgerald M, Maekawa AS, Kovacs CS, et al. Mutation of foxl1 Results in Reduced Cartilage Markers in a Zebrafish Model of Otosclerosis. Genes (Basel). 2022;13(7).

55. Drabkin M, Jean MM, Noy Y, Halperin D, Yogev Y, Wormser O, et al. SMARCA4 mutation causes human otosclerosis and a similar phenotype in mice. J Med Genet. 2024;61(2):117–24.

56. Fabbris C, Molteni G, Tommasi N, Marchioni D. Does pregnancy have an influence on otosclerosis? J Laryngol Otol. 2022;136(3):191–6.

57. Kovacs CS. Calcium and bone metabolism in pregnancy and lactation. J Clin Endocrinol Metab. 2001;86(6):2344–8.

58. Kovacs CS. Calcium and bone metabolism during pregnancy and lactation. J Mammary Gland Biol Neoplasia. 2005;10(2):105–18.

59. Kovacs CS. Maternal Mineral and Bone Metabolism During Pregnancy, Lactation, and Post-Weaning Recovery. Physiol Rev. 2016;96(2):449–547.

60. Mardinian K, Adashek JJ, Botta GP, Kato S, Kurzrock R. SMARCA4: implications of an altered chromatin-remodeling gene for cancer development and therapy. Molecular cancer therapeutics. 2021;20(12):2341–51.

61. Schoenfeld AJ, Bandlamudi C, Lavery JA, Montecalvo J, Namakydoust A, Rizvi H, et al. The genomic landscape of SMARCA4 alterations and associations with outcomes in patients with lung cancer. Clinical Cancer Research. 2020;26(21):5701–8.

62. Tian Y, Xu L, Li X, Li H, Zhao M. SMARCA4: Current status and future perspectives in non-small-cell lung cancer. Cancer letters. 2023;554:216022.

63. Chen A, Zhong L, Lv J. FOXL1 overexpression is associated with poor outcome in patients with glioma. Oncology Letters. 2019;18(1):751–7.

64. Vuorinen EM, Rajala NK, Ihalainen TO, Kallioniemi A. Depletion of nuclear import protein karyopherin alpha 7 (KPNA7) induces mitotic defects and deformation of nuclei in cancer cells. BMC cancer. 2018;18(1):325.

65. Laurila E, Vuorinen E, Savinainen K, Rauhala H, Kallioniemi A. KPNA7, a nuclear transport receptor, promotes malignant properties of pancreatic cancer cells in vitro. Exp Cell Res. 2014;322(1):159–67.

66. Diaz C, Thankam FG, Agrawal DK. Karyopherins in the Remodeling of Extracellular Matrix: Implications in Tendon Injury. J Orthop Sports Med. 2023;5(3):357–74.

67. Mauri G, Sartore-Bianchi A, Russo AG, Marsoni S, Bardelli A, Siena S. Early-onset colorectal cancer in young individuals. Molecular oncology. 2019;13(2):109–31.

68. Tavazzani E, Spaiardi P, Contini D, Sancini G, Russo G, Masetto S. Precision medicine: a new era for inner ear diseases. Front Pharmacol. 2024;15:1328460.

69. Miller SA, Dykes DD, Polesky HF. A simple salting out procedure for extracting DNA from human nucleated cells. Nucleic Acids Res. 1988;16(3):1215.

70. Abdelfatah N, Mostafa AA, French CR, Doucette LP, Penney C, Lucas MB, et al. A pathogenic deletion in Forkhead Box L1 (FOXL1) identifies the first otosclerosis (OTSC) gene. Human genetics. 2022;141(3):965–79.

71. Ziff JL, Crompton M, Powell HR, Lavy JA, Aldren CP, Steel KP, et al. Mutations and altered expression of SERPINF1 in patients with familial otosclerosis. Hum Mol Genet. 2016;25(12):2393–403.

72. Abecasis GR, Cherny SS, Cookson WO, Cardon LR. Merlin--rapid analysis of dense genetic maps using sparse gene flow trees. Nat Genet. 2002;30(1):97–101.

73. DePristo MA, Banks E, Poplin R, Garimella KV, Maguire JR, Hartl C, et al. A framework for variation discovery and genotyping using next-generation DNA sequencing data. Nat Genet. 2011;43(5):491–8.

74. Van der Auwera GA, Carneiro MO, Hartl C, Poplin R, Del Angel G, Levy-Moonshine A, et al. From FastQ data to high confidence variant calls: the Genome Analysis Toolkit best practices pipeline. Curr Protoc Bioinformatics. 2013;43(1110):11.0.1–.0.33.

75. Li H, Handsaker B, Wysoker A, Fennell T, Ruan J, Homer N, et al. The Sequence Alignment/Map format and SAMtools. Bioinformatics. 2009;25(16):2078–9.

76. Cingolani P, Platts A, Wang le L, Coon M, Nguyen T, Wang L, et al. A program for annotating and predicting the effects of single nucleotide polymorphisms, SnpEff: SNPs in the genome of Drosophila melanogaster strain w1118; iso-2; iso-3. Fly (Austin). 2012;6(2):80–92.

